# Sleep selectively and durably enhances real-world sequence memory

**DOI:** 10.1101/2024.01.10.575038

**Authors:** N.B Diamond, S. Simpson, D. Baena Pérez, B. Murray, S. Fogel, B. Levine

## Abstract

Sleep is thought to play a critical role in the retention of episodic memories. Yet it remains unclear whether and how sleep actively transforms memory for specific experiences. More generally, little is known about sleep’s effects on memory for multidimensional real-world experiences, both overnight and in the days to months that follow. In an exception to the law of forgetting, we showed that sleep actively and selectively improves retrieval of a one-time real-world experience (a controlled but immersive art tour) – specifically boosting memory for the order of tour items (sequential associations), but not perceptual details from the tour (featural associations). This above-baseline increase in sequence memory was not evident after a matched period of wakefulness. Moreover, the sleep-induced advantage of sequence over featural memory grew over time up to one-year post-encoding. Finally, overnight polysomnography showed that sleep-related memory enhancement was associated with the duration and neurophysiological hallmarks of slow-wave sleep previously linked to neural replay, particularly spindle-slow wave coupling. These results suggest that sleep serves a crucial and selective role in enhancing sequential organization in episodic memory at the expense of specific details, linking sleep-related neural mechanisms to the transformation and enhancement of memory for complex real-life experiences.

**Significance Statement:** Sleep affects the retention of episodic memories. Yet, it remains unclear whether sleep active transforms how we remember past experiences, overnight and beyond. We investigated memory for different dimensions underlying a dynamic real-world event – sequential associations versus atemporal featural associations – before and after sleep or wakefulness, and serially up to a year later. Sleep actively and selectively enhanced sequence memory, with this preferential sequence retention growing with time. Overnight memory enhancement is associated with the duration and neurophysiological hallmarks of slow-wave sleep previously linked to sequential neural replay, particularly spindle-slow wave coupling. Our findings support an active role for sleep in transforming different aspects of real-world memory, with sequence structure coming to dominate long-term memory for dynamic real-world experiences.

---

Episodic memory allows us to retrieve specific experiences from our past, mentally reactivating co-occurring perceptual details (e.g., the sight of syrup on my sister’s white dress), and replaying the sequence of events in which they were embedded (driving to the diner, and the graduation ceremony after breakfast (1). The law of forgetting (i.e., transience) dictates that these memories fade with time (2–4). Yet memory does not decline simply as a function of clock- and calendar-time (5). Forgetting depends, among other factors, on cognitive states intervening between encoding and retrieval – especially sleep. A century of experimental evidence demonstrates that sleeping after an event results in better retention than matched periods of wakefulness (6–8). It remains unclear, however, whether sleep’s effects on episodic memory derive from passively sheltering it from interference (7, 9, 10), or from actively transforming memories into more durable or adaptive formats (8, 11, 12). Does sleeping after a given experience make us merely forget it less, remember it better, or remember it *differently*?

Active systems consolidation theories are rooted in the observation of sleep-specific neural dynamics that are linked to the preservation of newly encoded experiences (8). Initially labile episodic memory traces are thought to be stabilized via repeated and time-compressed hippocampal-to-cortical replay during sleep (13, 14), presenting a mechanism by which one-time experiences can stick in memory: they endogenously recur, in some form, during sleep. Sleep-related replay occurs during hippocampal sharp-wave ripples, which in turn are hierarchically nested in sleep spindles (brief 11-16 Hz bursts) and slow waves (0.5 – 4 Hz oscillations) during slow-wave sleep (SWS) (15, 16). Evidence in humans is consistent with this proposed mechanism – episodic memory performance varies with SWS duration and microstructure (e.g., (17); for reviews, see (8, 18)), particularly temporal or phase-based spindle-slow wave coupling, which is thought to facilitate replay-linked synaptic plasticity (15, 19).

However, the evidence for sleep’s active role in shaping episodic memory is mixed, with failed replications and task-dependence rendering unclear which retrieval tasks or processes should be most affected (20, 21). Moreover, the presumption that sleep-related consolidation transforms memories from specific (i.e., episodic) to abstracted,

generalized knowledge via hippocampal-to-cortical information transfer (8, 15, 22), does not explain the role of sleep (if any) in stabilizing or enhancing episodic memory for specific experiences. Relatedly, if sleep adaptively transforms rather than passively preserves episodic memory, then some elements of memory should improve in absolute terms from pre- to post-sleep, beyond a mere reduction in forgetting relative to wakefulness. To our knowledge, however, there is little or no evidence for the predicted sleep-induced above-baseline enhancement of episodic memory, owing perhaps to insensitive behavioural measurement. Sleep may in some cases improve non-episodic memory tasks like motor sequence learning or statistical learning (23, 24) (but see (25)), but in studies of episodic memory, sleep at best reduces the amount of forgetting relative to wakefulness (18) (but see (26)).

These discrepancies may be attributable in part to the differential forgetting of heterogenous episodic representations over time (27–30), and perhaps sleep. The two defining types of associations underlying episodic memories, linked to distinct hippocampal-cortical networks, are i) associations linking simultaneously encoded event features (e.g. an object and its colour), and ii) sequential associations bridging temporal gaps between successive events (e.g., what happened at time *t* and time *t+1*) (1, 31–33). Sleep studies are often equivocal with respect to which aspects of episodic memory ought to be most affected by sleep, testing memory for laboratory stimuli in binary fashion (recalled or not). However, the mechanics of sleep-related replay suggest that it should preferentially enhance sequential organization in memory representations – associations bridging experiences across time and space – as opposed to simultaneously encoded features. In rodents, neural replay recapitulates the spatiotemporal order of waking experience, and compresses behavioural-timescale gaps between encoded events (i.e., seconds or longer) down to the millisecond-level timescale of synaptic plasticity, supporting subsequent post-sleep navigation (13, 34).

Accordingly, in humans, sleep benefits the detection of temporal rules or regularities more than other kinds of associative relations across studies (35), and has also been found to reduce forgetting of temporal order information relative to wakefulness (36, 37). Yet sleep’s particular influence on retrieval of sequential versus atemporal associations remains unknown. Furthermore, little is known about sleep’s influence on real-world episodic memory – which is more durable than memory for laboratory stimuli (38) – beyond delays of a few hours or a day, despite evidence that consolidation or transformation unfolds over weeks to months or longer (5, 39, 40).

The present study examined (i) how memory for sequential versus featural associations from a single real-world episode transforms with sleep and over longer timescales (Study 1), (ii) whether sleep *per se* is necessary for differential sequence versus featural memory transformation (Study 2), and (iii) whether sleep-related memory transformation is related to the duration and predicted neural mechanisms of slow-wave sleep (Study 3). We devised a novel test of episodic memory for a controlled but immersive real-world event – the ‘Baycrest Tour’ (31, 38), an audio-guided walking tour of artwork, paralleling the goal-directed and ecologically-scaled paradigms used in rodent research (see Figure 1). Trial-unique and difficulty-matched memory tests probed both the sequence structure (e.g., “You encountered the bicycle sculpture before the scale model of Baycrest”) and static perceptual features (e.g., “The bicycle sculpture was made of metal”) of the tour items before and after a night of sleep, and at subsequent delays from days to over a year after the tour. We hypothesized that sleep transforms episodic memory across short- and long-term timescales by preferentially enhancing sequential structure in memory over atemporal perceptual associations, and that this enhancement would be related to SWS duration and its physiological hallmarks: spindles, slow waves, and particularly spindle-slow wave coupling.

**Figure 1.**
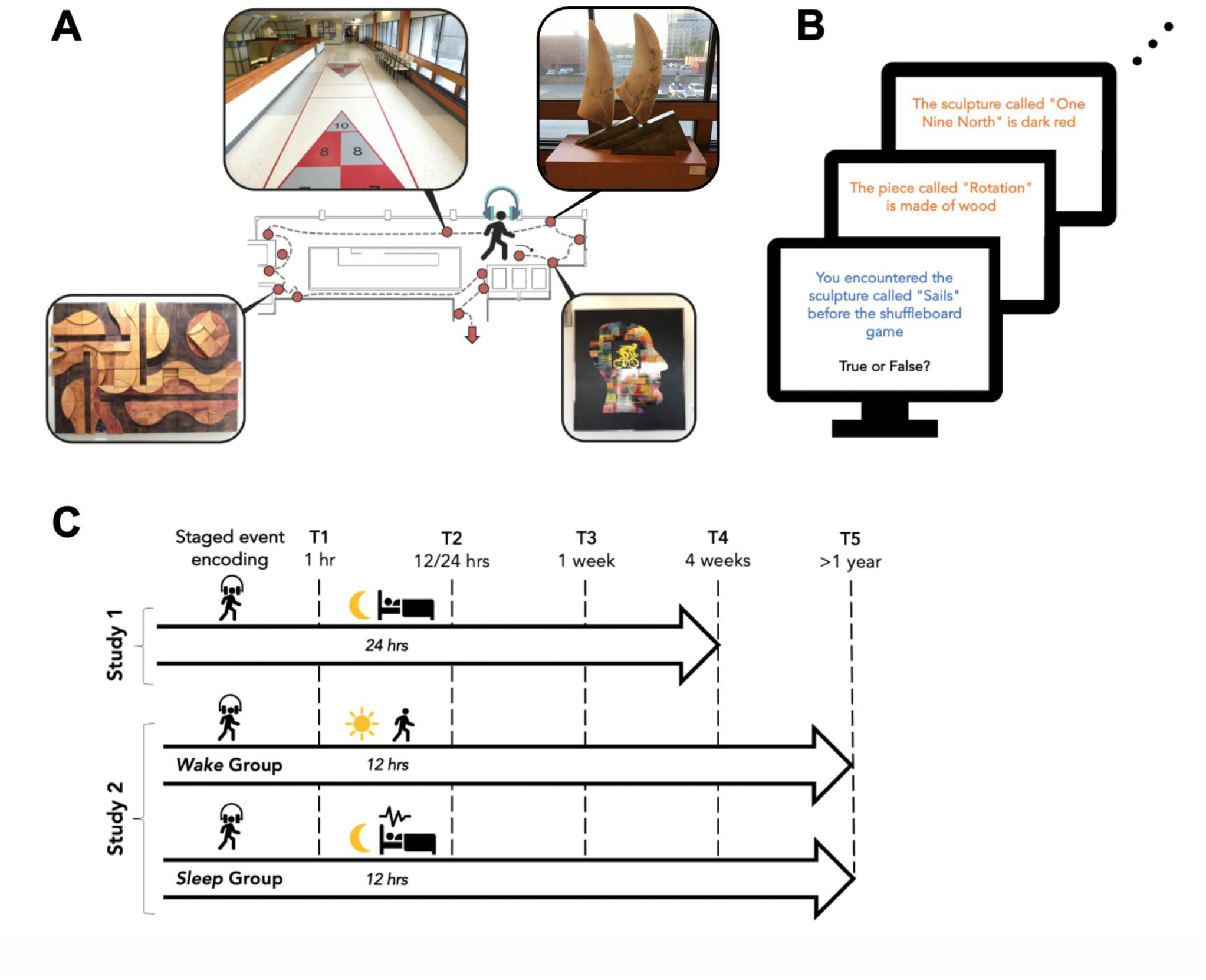
***A.*** Depiction of the beginning of the audio-guided tour route (dashed line), including the locations and select examples of the first 13 target events. ***B.*** Each memory test included trial-unique, difficulty-matched questions probing sequence (e.g., *You encountered the sculpture called “Sails” before the shuffleboard game; blue text*) and featural (e.g., *The piece called “Rotation” is made of wood; orange text*) information from the Tour. ***C.*** Study design: Four unique true/false recognition memory tests were administered at four time points (T1-T4), with test order counterbalanced across participants. In **Study 1** *(n* = 53), the tour was completed during regular business hours, with T1 occurring at 1 hour and T2 occurring at 24 hours post-encoding. **Study 2** participants underwent identical encoding conditions but were randomized to Sleep (*n* = 39) or Wake (N = 38) groups, with T1 occurring at 1 hour post-encoding and T2 occurring after 12 hours in both groups. Critically, participants either slept overnight in the sleep laboratory (Sleep group) or stayed awake (Wake group) during this 12-hour delay. T3 and T4 occurred at 1 week and 1 month post-encoding for both Study 1 and Study 2. Fifty-six Study 2 participants (*n* = 33 Sleep, *n* = 23 Wake) completed a post-hoc fifth test (T5) 15 months post-encoding. **Study 3** (not depicted) comprised the Study 2 Sleep group plus additional recruits to the sleep laboratory between T1 and T2 (total *n* = 49) to increase power for individual differences analyses linking sleep neurophysiology to memory change.

In Study 1, we found differential transformation of sequence versus featural memory from one hour to one month: sequence memory significantly *increased* above baseline over a 24-hour delay including a night of sleep, and remained stable up to a month later, whereas featural memory declined monotonically at the classic Ebbinghausian forgetting rate (linearly on an approximately logarithmic timescale) (3, 4). In Study 2, we replicated Study 1 and demonstrated that sleep (contrasted with a matched period of wakefulness in a between-subjects design) was necessary for the boost in sequence memory, also finding that the sleep-related enhancement of sequence memory held at a 15-month follow-up. In Study 3, using overnight polysomnography recording, we found that individual differences in slow-wave sleep duration and neurophysiology – particularly spindles, slow waves, and their coupling – were positively associated with enhanced memory for this real-life event.

## Results

### Sleep-related enhancement of sequence versus featural memory

As expected, given the law of forgetting, overall memory accuracy declined from T1-T4 (*χ*^2^(1) = 68.54, *p* < .001; Figure 2A), but this effect was qualified by a crossover interaction between Time and Retrieval Type (sequence, featural) (*χ*^2^(1) = 53.77, *p* < .001). The significant advantage for featural memory over sequence memory observed just after encoding (T1: *z* = 3.26, *p* = .001, *d* = .46) flipped in direction after a night’s sleep (T2: *z* = .27, *p* = .789, *d* = .12), with a significant advantage for sequence over featural memory at one week (T3: *z* = 2.26, *p* = .024, *d* = .43) that widened over time (T4: *z* = -3.65, *p* < .001, *d* = .83). Sequence and featural memory accuracy did not differ overall (*χ*^2^(1) = .69, *p* = .405). Considering the overnight trajectories of sequence versus featural memory: sequence memory increased significantly overnight (*z* = 2.65, *p* = .008, *d* = .33), whereas featural memory significantly declined (*z* = -2.38, *p* = .018, *d* = -.29) (Figure 2B).

**Figure 2.**
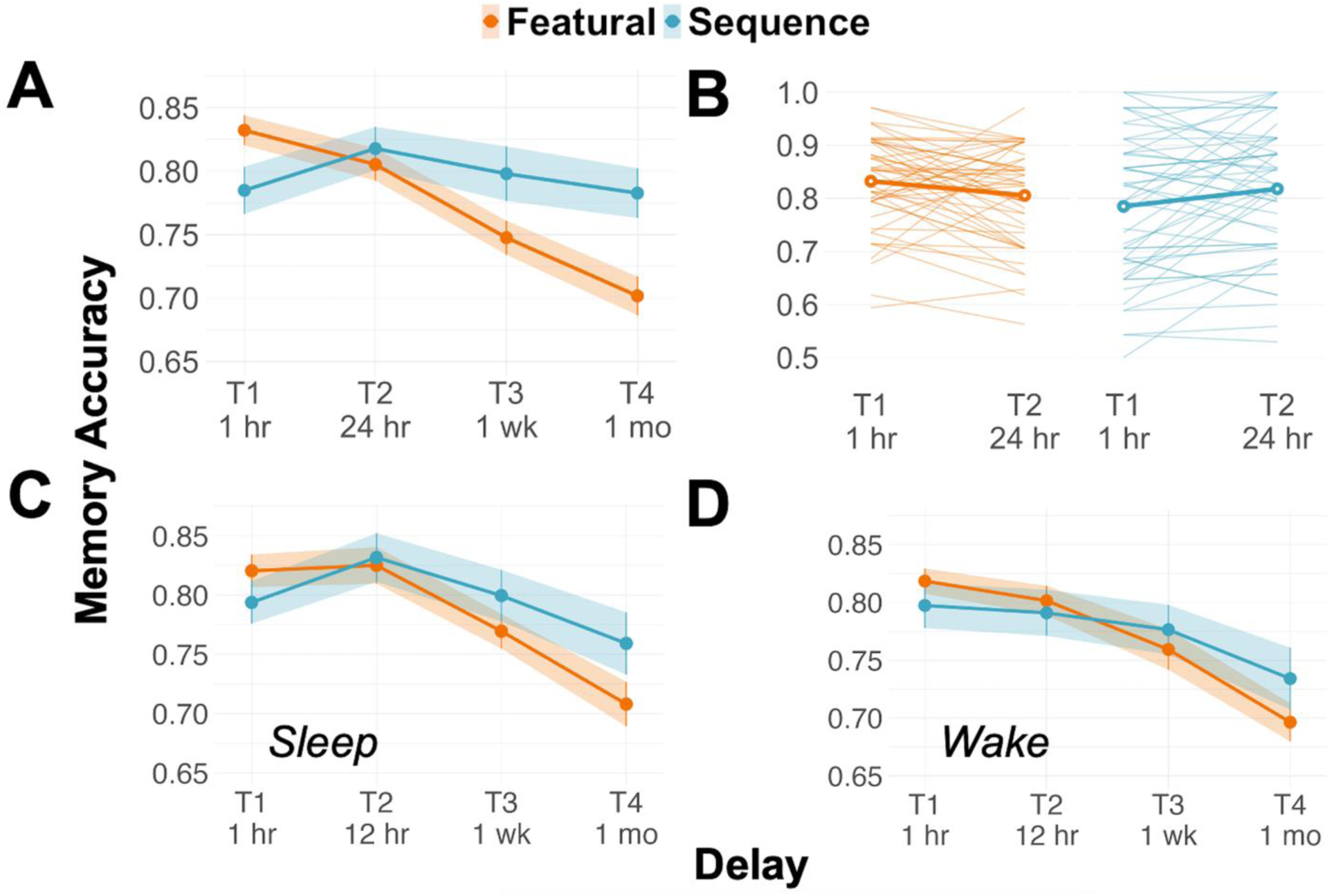
Sequence (blue) versus featural (orange) memory accuracy across T1-T4, approximately logarithmically spaced across time to fit a negatively-accelerated forgetting function. Dots depict means, vertical lines and shaded regions depict between-subjects standard errors. ***A.*** Study 1. ***B.*** Accuracy change from T1-T2 in Study 1: thin lines depict individual Study 1 participants’ memory, and circles and thick lines depict group averages; sequence change and featural change were not significantly correlated (*r*(51) = .002, *p* = .986). ***C.*** Study 2 *Sleep Group,* who slept in the sleep lab between T1 and T2. ***D.*** Study 2 *Wake Group,* who did not sleep between T1 and T2.

**Figure 3.**
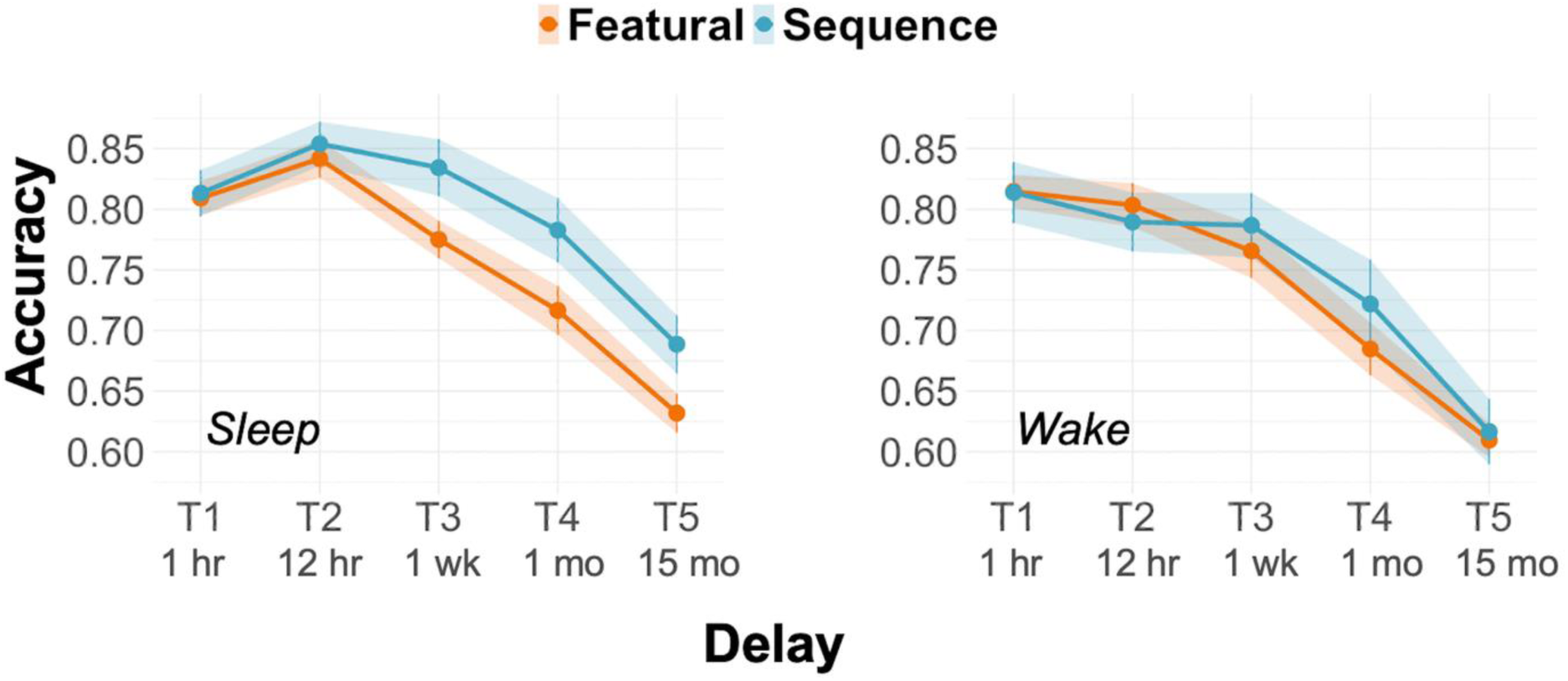
Sequence versus featural memory accuracy from T1 to T5 in the Sleep and Wake groups. The sleep-related advantage for sequence over featural memory performance was held at 15 months post-encoding.

The preferential effect of sleep on sequence versus featural memory could not be explained by item characteristics such as difficulty or true/false phrasing (see Supplementary Figure S1). Sequence memory improved as the inter-item lag (i.e. spatiotemporal distance) between the tour events in each pair increased in both Study 1 and 2 (see Supplementary Figure S2), replicating prior sequence memory assessments (41, 42), and this sequence lag effect did not vary with time – even fine-grained sequence memory accuracy (for item pairs that were adjacent or had only one intervening item) was retained from an hour (T1) to a month (T4) after encoding.

Study 2 participants were randomly assigned to either a Wake or Sleep condition (see Figure 1). Here, we first replicated the findings observed in Study 1 in the Sleep condition, where a main effect of Time (T1 – T4) (*χ*^2^(1) = 60.40, p < .001) was once again qualified by a Time × Retrieval Type crossover interaction (*χ*^2^(1) = 13.85, p < 0.001) (Figure 2C). Sequence memory performance exceeded featural memory performance after one month (T4; *z* = 2.11, *p* = 0.035, *d* = 0.47), but not earlier (*z’s* < |.17|, *p’s* > .097) (Figure 2C). As in Study 1, sequence memory significantly increased overnight (*z* = 2.76, *p* = .006, *d* = 0.47), but featural memory did not (*z* = -0.43, *p* = .670, *d* = -0.04; see Figure S3 for participant-level data).

Considering the Wake group, sequence and featural memory did not differ at any time points (*z*’s < |1.5|, *p*’s > .143), nor was there a T1-to-T2 increase in sequence memory as seen in Study 1 and the Study 2 Sleep group; indeed, sequence memory declined, albeit non-significantly (*z* = -.45, *p* = .655, *d* = -0.06). When the Sleep and Wake groups were analyzed in the same model (T1-T2), there was a significant Group X Time interaction (*χ*2(1) = 5.86, *p* = .016), such that overall memory accuracy was greater after a period of Sleep (*z* = 2.29, *p* = .022, *d* = 0.315) but not Wake (*z* = -1.13, *p* = .258, *d* = -0.213), although the Group x Time x Retrieval Type interaction was not significant (*χ*2(1) = 0.34, *p* = .558). Memory performance differences across the Sleep and Wake groups could not be attributed to time-of-day differences as their scores were nearly identical at T1 (see Figure 2); nor were there group differences in chronotype or self-reported sleep habits (see *Supplementary Methods*). A model examining all four time points revealed the expected main effect of Time (*χ*2(1) = 78.62, *p* < .001) and the Time X Retrieval Type interaction (*χ*2(1) = 9.16, *p* = .002), but no significant interactions with Group (Group X Time: *χ*2(1) = 0.39, *p* = .533; three-way interaction: *χ*2(1) = 0.34, *p* = .560).

### Long endurance of sleep-related mnemonic advantage at 15 months post-encoding

Having found relatively strong sequence memory retention at the longest delay in our original design (1 month), we created a fifth memory test (matching the properties of the first four and drawing test items from them equally; see Methods), delivered to Study 2 participants (33 and 23 from the Sleep and Wake groups, respectively) after approximately 15 months, roughly approximating the next proportionately-spaced timepoint given a negatively-accelerated forgetting function across time. As expected, overall memory performance declined over time (*χ*^2^(1) = 169.72, *p* < .001), but it remained above chance at T5, *M* = 0.64, *SD* = 0.03). The mnemonic enhancement conferred by one night of sleep (versus wake) held at 15 months post-encoding (Group × Time [T1, T5] interaction; *χ*^2^(1) = 4.46, *p* = .035). Although the three-way Group × Time × Retrieval Type interaction was not significant (*χ*^2^(1) = 0.95, *p* = .331), hypothesis-driven planned comparisons confirmed sequence memory performance from T1 to T5 was preserved by sleep relative to wake (*z* = -2.17, *p* = .030, *d* = -0.652); there were no group differences in T1-T5 change in featural memory performance (*z* = -0.81, p = .417, *d* = -0.350). While the Study 2 long-term (T5) follow-up participants were composed of a subset of the full sample, they were demographically similar to the main sample (see Supplementary Materials) and there was no *a priori* reason to expect that these individuals would show a disproportionate advantage for sequence over featural memory.

### Overnight memory transformation is linked to sleep neurophysiology

Study 3 participants – including the Sleep group from Study 2 and additional participants recruited for individual differences analyses – underwent overnight polysomnography recordings in the sleep laboratory between T1 and T2. When modelling sleep macrostructure, SWS duration was positively and selectively associated with overall memory enhancement (SWS duration × Time [T1, T2] interaction, *F*(1, 131) = 4.51, *p* = .036, partial eta squared = 0.03; see *Supplementary Figure S4, Table S3* for null effects of other sleep stages, and null interactions with Retrieval Type). This effect extended to SWS microstructure – both slow waves (half waves) and spindles during SWS were associated with overall memory enhancement, with both interacting with Time (*F*’s(1, 141) = 7.43, 8.66, *p*’s = .007, 004, partial etas squared = 0.05, 0.06; see Supplementary Table S4 and Figure S4). Critically, spindles coupled to slow waves – *but not uncoupled spindles* – were associated with overall memory enhancement (interactions with Time: *F*(1, 138) = 9.32, *p* = .003, eta squared = 0.06; *F*(1, 138) = 0.070, *p* = 0.792, eta squared ∼ 0.00, for coupled and uncoupled spindles respectively; see Supplementary Table S5 and Figure S5).

Given our prior results concerning the behavioural dissociation of sequence and featural memory change overnight, these measures are visualized separately in relation to coupled and uncoupled spindles (Figure 4). Overnight sequence memory enhancement was uniquely associated with coupled spindles, and significantly more so than uncoupled spindles. Featural memory showed a similar pattern, although the correlations between coupled versus uncoupled spindles were not significantly different (see Figure 4).

**Figure 4.**
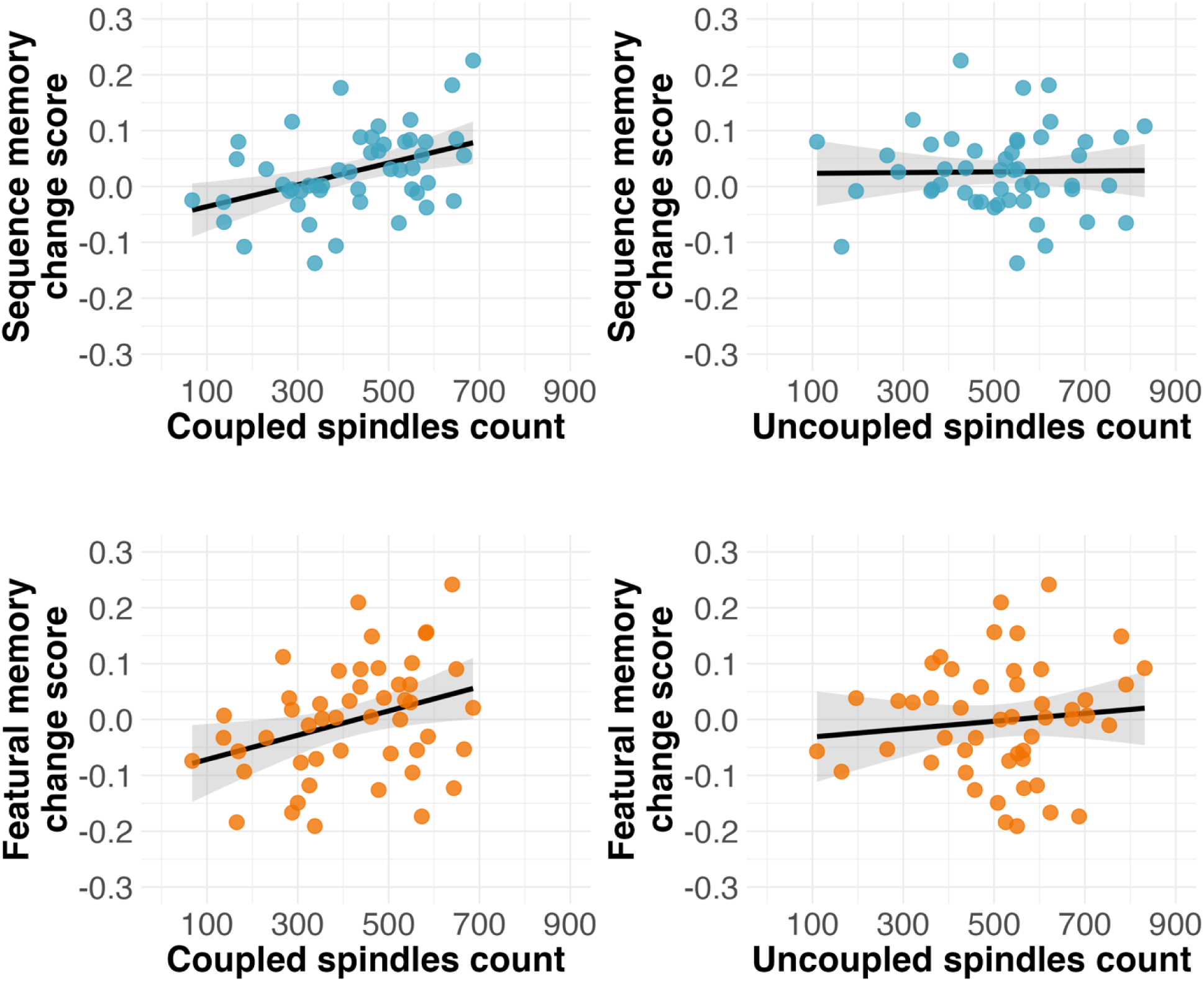
Sequence memory change (T2 – T1) is significantly associated with coupled (*r* (47) = 0.42, *p* = .003) but not uncoupled spindles (*r* (47) = 0.01, *p* = .921) in SWS. These correlations are statistically different from each other (Steiger’s *z* = 2.24, *p* = .029). Similarly, featural memory change (T2 – T1) is related to coupled (*r* (47) = 0.34, *p* = .018) but not uncoupled spindles (*r* (47) = 0.11, *p* = .451) in SWS. These correlations are not statistically different from each other (Steiger’s *z* = 1.21, *p* = .231).

## General Discussion

Memories fade with time, yet memory change is not uniform – while most details from our experiences are forgotten, some can persist with fidelity even after years (28). Sleep’s influence on the fate of our memories has been studied for at least a century. Theories of sleep-related memory consolidation hinge on the notion that sleep’s effects on memory are active and selective, leading memories to be qualitatively transformed, not merely protected (8, 17, 18). However, the case for preferential consolidation of certain types of episodic memory (e.g., emotional memoranda) over others has been complicated in recent years by failed replications (6, 20), and the effect of sleep on memory consolidation is still actively debated (10, 21). Using an immersive real-world encoding event, we found in two independent samples that sleep selectively and actively enhanced memory for sequence associations – but not featural associations – despite sequence and featural memory probes being matched on difficulty overall. This preferential enhancement of sequence memory was eliminated after a matched period of wakefulness. Lasting sleep-related enhancement of sequence over featural memory was evident across two independent samples and persisted after 15 months, extending previous findings of long-lasting sleep-related mnemonic enhancement (43) from laboratory stimuli to real-world experiences, specifically for sequential (versus static featural) associations.

In accordance with active, as opposed to passive or ‘permissive’ theories of sleep-dependent transformation or consolidation (12, 15), we found that the duration of SWS (but not other sleep stages) and its neural hallmarks, including slow waves and spindles, were associated with memory for a real-world experience. Crucially, spindles coupled to slow waves uniquely predicted overnight memory enhancement, aligning with contemporary models of hippocampal-neocortical communication in sleep-related mnemonic processing (13, 19) (for review, see (15)). Contrary to our hypothesis and behavioural evidence for sleep’s selective effect on sequence memory, SWS duration and neurophysiology predicted *both* sequence and featural memory change – although there was evidence for greater specificity to spindle-slow oscillation coupling for sequence relative to featural memory. We speculate that design differences mitigated the dissociation of sequence and featural memory as assessed at T2 in Studies 2 and 3.

In Study 1 there was a crossover interaction such that sequence memory improved in absolute terms whereas featural memory declined overnight. While the sleep effect on sequence memory replicated (and was in fact stronger) in Study 2, featural memory performance stayed flat from T1 to T2, after which it declined as expected across subsequent assessments. This unexpected overnight protection of featural memory may be attributable to the fact that Study 2 participants slept shortly after encoding (i.e., in less than 2 hours versus up to 15 hours in Study 1), suggesting that sleep may play a passive, interference-reducing role in protecting atemporal associations in contrast to an active strengthening of sequential associations. Accordingly, the interactions between Retrieval Type and Group (Study 2; Sleep versus Wake) and neurophysiology (Study 3) were attenuated; effects were instead observed for overall memory enhancement. Notably, sequence and featural forgetting functions crossed after a night’s sleep in all three experimental groups (between T1 and T2 in Study 1 and Study 2 Wake, and between T2 and T3 in Study 2 Sleep). Overall, the enhancement of sequence over featural memory attributable to a single night of post-encoding sleep – and linked to spindle-slow wave coupling – emerged as featural memory declined in an Ebbinghausian manner with time, and this enhancement remained robust after 15 months. These findings highlight the importance of pairing serial, independent, and delayed assessments in studies of human memory transformation.

This is to our knowledge the first evidence for a sleep-specific, above-baseline improvement in episodic memory retrieval, demonstrating an exception to the lawful decline of episodic memory with time (3, 4). This finding is also consistent with the hypothesized role of sequential and time-compressed replay, identified mainly in rodents (13) and more recently in humans (44), in the stabilization or enhancement of sequential organization in episodic memory (15, 35). Whereas real-life episodic memory encoding usually occurs through volitional exploration in large-scale environments, participants in most human memory studies passively encode discrete memoranda at a fixed location (e.g., words or pictures on a computer screen). In rodents, active self-motion through space (versus being passively transported) is necessary to facilitate subsequent sleep-related replay (34), and active real-world encoding improves human episodic memory relative to passive laboratory conditions (38). Recall of such action sequences encoded in large-scale space may therefore be enhanced by hippocampal replay during sleep to a greater degree than recall of simultaneously encoded perceptual associations and could rely on strengthening of different heteroassociative versus autoassociative circuits (45) or hippocampal-cortical networks (32). Active and immersive encoding paradigms such as the one used in this study may also explain the striking long-term retention of sequential structure in prior real-world episodic memory studies (31, 46, 47).

The differentiated long-term forgetting curves for sequence and featural memory have implications for theories of memory consolidation. First, this dissociation builds on decades-old findings that different elements of complex events are forgotten at different rates (30, 48), with associations across events being better remembered than associations within events (27). Systems consolidation theories, however, focus on sleep’s role in extracting gist or structure shared across similar repeated episodes (thought to be cortex-dependent), rather than preserving or enhancing episode-specific information (thought to be hippocampus-dependent (15, 22)). To the extent that sequential (i.e. spatiotemporal) information reflects an underlying event structure, the present results advance these theories, but also highlight that such structure can be extracted even from a one-shot event, much as flexible neural replay can emerge after a single maze traversal in rats (49). The persistence of sequence memory for a single experience up to a year raises the question of whether it comes to rely on cortical mechanisms, or rather reflects an enduring role for the hippocampus in remote episodic memory access and sequential organization (39, 50).

Why, then, is sequence information so sticky? Memories for the sequence in which events occurred provide the building blocks for future predictions and simulations (51), and enable retroactive and delayed linking of actions to their eventual outcomes, consistent with replay’s putative role in back-propagating reward value to preceding behavioural trajectories (52–54). Sequential associations might therefore form a latent structural representation – or cognitive map – upon which other episodic features (e.g., visual, auditory, tactile features) are laid (50, 55). Indeed, spatiotemporal associations between events automatically shape the way we search memory during natural, unconstrained free recall (31, 56), and sleep tends to benefit free recall tasks more than cued recall or recognition (6). This might explain why spatiotemporal aspects of episodic memory are privileged over atemporal featural associations after sleep and in the weeks and months that follow.

Marrying a controlled real-world encoding paradigm, between-subjects manipulation of sleep, serial assessment of different components of episodic memory with matched trial-unique tests, and individual differences in overnight electrophysiology, this study deepens our understanding of how sleep transforms memory over the short and long term. The observation of enduring overnight gains in sequence memory for a one-shot event, linked to underlying neural mechanisms of slow-wave sleep, provides critical new evidence for theories of memory consolidation in humans.

## General methods

### The Baycrest Tour

Participants underwent a 20-minute audio-guided walking tour of a section of Baycrest Health Sciences Centre lined with artworks (Figure 1a; see also (31)). The audio guide was presented on a portable digital player with headphones, similar to those used in museums. Thirty-three tracks corresponding to sequential items along the tour route were presented in a fixed order. Each track provided information about the item (e.g., the artist’s name, the medium), instructed participants to examine the item (followed by 10 seconds of silence), and then directed participants to walk to the next item, whereupon the participant initiated the next track. The guide thus controlled – in addition to featural content and sequence – the encoding duration for each item, allowing for individual differences in walking speed between items. The tour took 20.83 minutes on average (range of tour duration = 16-28 minutes across studies). The audio guide is openly accessible at our OSF repository: https://osf.io/bxm5w/.

### Memory test design

All memory tests were implemented using the Qualtrics online platform. 276 true/false statements were created that pertained to either features (details) or sequences. Featural statements refer to details such as the colour or shape of a given piece (e.g., “The sculpture called One Nine North is dark red”), whereas sequence statements refer to the spatiotemporal order of pairs of items from the tour (e.g., “You encountered the sculpture called One Nine North before the Spiro Family Gardens painting”). Sequence statements pertained to 25 ‘target events’ encoded during the tour in a fixed, ordinal position, as ensured by the audio guide and the unidirectional track-like layout of the tour. We use the word ‘sequence’ to encompass temporal and spatial associations, which are inherently confounded in unidirectional real-world navigation (50). Sequence statements were sub-classified as Near, Medium, or Far based on their encoded inter-item lag (i.e., the number of intervening target items between the pair in question; 0-1, 2-3, or 4-6), which was taken into consideration when constructing the test forms. False statements were created by altering details or reversing the sequence. See *Supplementary* for a more complete description of test construction.

The statements were distributed across four equivalent 69-item test forms (considering proportions of true/false, featural/sequence statements, sequence lag, reference to different target events across the tour, and item difficulty based on pilot testing at a one-day delay in an independent sample). The order of test forms across test sessions T1 - T4 was counterbalanced using a Latin square design, creating four different form orders (ABDC, BCAD, CDBA, DACB) to which participants were randomly assigned, minimizing any order effects in group-level results.

Given the generally high performance at the fourth time point (one month), we created a fifth recognition memory test (T5) *post hoc* to probe the memory performance at an even longer delay that is typical of naturalistic autobiographical memory research but rarely probed with a controlled and verifiable laboratory assessment – approximately 15 months (Study 2). As this was not part of the original test design, 100 items (50 featural and 50 sequence) were selected from the original four test forms in equal proportion for the T5 test; 49 featural and 42 sequence remained after exclusion due to a coding error that resulted in duplicate statements (8 items) or poor average performance (1 item).

The featural and sequence item groups administered at T5 were matched for difficulty (based on T1-T4 item analysis; see *Supplementary*). T5 items also sampled all tour target events/placements and inter-item lag (for sequence items) equally. Although these items were repeated from earlier tests, the effects of repetition were expected to be minimal given the passage of time. In any case, as the featural and sequence items were matched for superficial characteristics of prior presentation, any differences between the two item types could not be attributed to these factors.

### Procedure

Participants were recruited via the Rotman Research Institute participant database or from online advertisements. Participants were screened for neurological or psychiatric disorders or other illnesses affecting cognition, substance abuse, or medications affecting memory or sleep (e.g., benzodiazepines), and prior exposure to the tour location. Study 2 and 3 participants were asked to refrain from ingesting alcohol and other psychoactive drugs the day prior to and the day of the experimental session.

Caffeinated beverages were permitted for all habitual coffee/tea drinkers on the day of the experiment. Study 2 participants randomized to the Wake group agreed not to nap during the day.

Both Study 1 and 2 used our Baycrest Tour 2.0 encoding paradigm (see also (31)) distinct in content and location from Baycrest Tour 1.0 used in (28, 38). Participants entered Baycrest on the first floor such that the tour location (second floor) was avoided. Following instructions and practice with the digital audio players in a private testing room, they were escorted via elevator to the tour start position. They completed the tour independently, with an experimenter unobtrusively following to ensure adherence to the protocol and to address potential technical issues. Following the tour, they returned to the testing room and completed a roughly 45-minute battery of questionnaires and neuropsychological tests (see *Supplementary*), followed by Test 1, administered online (via Qualtrics.com) to ensure consistency with Tests 2-4.

In Study 1, we assessed feature versus sequence memory for the Baycrest Tour across four test sessions, with sleep occurring between T1 and T2. The tour and first testing session were conducted during normal business hours according to the participants’ availability, with T2 completed online 24 hours later.

Study 2 participants consented to random assignment to Sleep or Wake groups at the time of recruitment; randomization occurred after study enrollment to reduce bias in recruitment into the Sleep or Wake condition should someone prefer to be enrolled in one condition over the other (see Figure 1B). The Sleep group completed the Baycrest tour at 6:30 PM and was subsequently escorted via taxi to Sunnybrook Hospital (a 15-minute drive from Baycrest) where the sleep lab is located. The sleep window was between 10:30 PM and 6:30 AM followed by T2 within 45 m of waking. Participants in the Wake group completed the tour between 9:00 - 10:00 AM and were instructed to stay awake for the next 12 hours before T2. As in Study 1, Study 2 participants completed T1 testing in the laboratory and T2-T4 (and T5) remotely. All memory tests were administered on Qualtrics.com.

After randomization to Sleep or Wake groups in Study 2 was completed, we continued to recruit participants into the Sleep group to increase sample size for the assessment of brain-behaviour relationships derived from sleep EEG (Study 3). This phase of the study was terminated after 10 participants were tested due to the onset of the COVID-19 pandemic.

Tests 3-4 were scheduled at one week and 1 month post-encoding for all studies, with the ancillary T5 testing occurring 15 months post-encoding (Study 2). The increasing delay across test intervals approximates the well-established power function of forgetting (4). The classic negatively accelerated forgetting function (3, 4) produces a linear decline across the roughly logarithmic test spacing (we note that this function was established in research that did not distinguish among elements of episodic memory, such as features and sequences.). Participants received scheduled email reminders in advance of T2-T5 testing, including instructions to complete the tests within the required time window. Tests completed at delays within 15% of the target time (relative to the total time elapsed from encoding) were accepted (see *Supplementary Table S1* for testing lags and exclusions).

### Polysomnography

Conventional polysomnography (PSG) data was collected from Study 2 and Study 3 participants using the Compumedics system (Australia) in accordance with American Academy of Sleep Medicine standards with frontal (F3, F4), central (C3, C4), and occipital (O1, O2) placement following the international 10-20 system, referenced to the contralateral mastoid (A1, A2; see *PSG Methods in Supplementary*). Sleep scoring was carried out and reviewed by registered sleep technologists at the Sunnybrook Health Sciences Centre. Movement and arousal artifact identification was conducted manually by expert raters. Spindles detected from sites where they appear maximally, at C3 and C4 derivations during artifact-free N2 and N3 sleep using a well-established, automated and validated method (57) using the EEGlab plugin (58) “*detect_spindles2.2*” written for MATLAB R2019a (The MathWorks Inc.) (see *Supplementary Polysomnography Methods* for more details). Consideration of fast and slow spindles separately did not add new information to hypothesis-driven analyses. Half waves (a measure of slow wave activity) were extracted from artifact-free N2 and N3 sleep from the sites where they appear maximally, at F3 and F4 derivations using a well-established automatic period amplitude analysis (PAA) method using the EEGlab “*PAA*” plugin, written for MATLAB R2019a (The MathWorks Inc., Natick, MA, United States). Spindles and slow waves were visually verified following automated detection (see *Supplementary Polysomnography Methods* for a more detailed description). Please see Supplemental Material for a full list of polysomnography descriptive data (*Supplementary Table S2*).

### Slow Wave-Spindle Coupling

Using the slow wave negative peak latencies from F3 and F4 and the spindle peak latencies from C3 and C4, we performed coupling detection procedures using the approach originally developed and validated by Mölle and colleagues (16, 59–63) employing EEGlab-compatible software (58) written for MATLAB R2019b (The MathWorks Inc.). To ensure that spindles would only be coupled to slow waves detected on the same hemisphere, electrode pairings F3-C3 and F4-C4 were made. This procedure involved building a time window of 4 seconds around the negative peaks of the detected slow waves. The spindles were identified as coupled slow wave-spindle (SW-SP) complexes if the spindle peak latency fell within a 4-second time window of the slow wave peak detected in the corresponding slow wave channel. Alternatively, spindles were marked as uncoupled if the spindle peak latency occurred outside the 4-second time window. The detection results for each electrode pairing were averaged and then used in the final analyses. To address duplicate detection of the same slow wave event by the different channels, we removed events in which the latency difference was lower than the minimum duration threshold for the half wave (0.125 seconds). Lag was measured as the distance between the slow-wave negative peak and the spindle onset. The lag variance was then calculated for each individual as a measure of coupling strength.

Additionally, the phase of the bandpass-filtered slow-wave signal in radians at the spindle peak latency was computed (see *Supplementary Figure S6*). The mean direction of the phase angles for all coupled spindle events was determined using the CircStat toolbox (64). Hilbert transform was applied to extract the preferred phase of SW-SP coupling for each participant averaging all individual events’ preferred phases.

Then, we perform uniformity tests (Rayleigh test) using positive slow wave peaks as the predefined mean direction (V-test). For the purposes of this paper, we analyzed the count of coupled and uncoupled spindles in SWS averaged across channels.

### Analysis

All analyses were conducted in RStudio version v1.4.1106 (R Core Team, 2019). For our main behavioural models, we used generalized linear mixed effects models (GLMMs) with a logit link function, using the *glmer* function from the *lme4* package in R. This allowed us to predict accuracy (correct versus incorrect) on a trial-wise basis from both Time (T1-T2 or T1-T4) and Retrieval Type (featural versus sequence; contrast-coded). For omnibus effects, we modelled Time as a continuous variable (mean-centred), based on our theoretically-motivated test spacing, as described above. GLMMs included a random intercept for subject and test item. Study 2 analyses included an additional fixed effect for Group (Wake, Sleep).

Omnibus significance tests were computed using Type III Wald chi-square tests, implemented using the *car* package in R. To decompose interactions into simple effects of Time or Retrieval type, we then re-ran models with Time as a categorical variable and reported pairwise difference model-fitted estimated marginal means (using the *emmeans* package in R). We report z-scores and uncorrected *p*-values for these tests of estimated marginal means and also report Cohen’s *d* (on ‘raw’ and not model-fitted comparisons for interpretability; with paired or unpaired comparisons as appropriate, where paired effect sizes were computed as *M*_difference_/*SD*_difference_). For Study 3, linear effects models were used to test the effects of sleep macrostructure (N1, N2, N3, and REM, with total sleep time (TST) as a covariate of no interest) on memory (effects on memory change were modelled by interacting each PSG predictor with a Time (T1, T2) factor). We subsequently probed the putative key microstructure measures in SWS, spindles, and half wave counts averaged over central and frontal electrodes, respectively on memory performance. A separate analysis was run to test the effects of coupled versus uncoupled spindles on memory performance. For all models, Time (T1, T2) and Retrieval Type (sequence, featural) were treated as fixed-effects and participants were treated as a random effect. Partial eta squared values are reported as measures of effect size for linear mixed effects models (using the *effectsize* package in R). Follow-up tests of the association between sleep neurophysiology parameters and memory change scores (T2 – T1 scores) were assessed with Pearson’s product-moment correlations using the *cor.test* function in R. Data analysis code and data are openly available in our OSF repository: https://osf.io/bxm5w/. Additional statistical analysis details are reported in the Supplementary Analyses.

## Acknowledgements

We thank Yarden Levy, Laryssa Levesque, Alissa Papadopoulos, Yushu Wang, Catherine Le, and Karla Machlab for technical assistance with designing Baycrest Tour 2.0 and developing the memory tests, Dana Jewell, Lily Lu and Arnel Panopio for their technical assistance with overnight sleep testing, Malcolm Binns for his analytical expertise, and Laura Ray for her support and guidance with the polysomnography analysis.

This project was funded by the Canadian Institutes of Health Research (MOP-148940, awarded to B.L.). N.B.D. was supported by an Ontario Graduate Scholarship. S.S. was supported by a Natural Sciences and Engineering Research Council of Canada Postgraduate Scholarship – Doctoral (PGS-D) and Early Professionals & Inspired Careers in AgeTech (EPIC-AT) Fellowship.

## Author Contributions

N.B.D. and S.S. made equal contributions to study conception and design along with the acquisition, analysis, and interpretation of data, and writing of this manuscript.

D.B. contributed to the analysis of PSG data and manuscript revision.

B.M. contributed to study design, along with data acquisition and interpretation, and manuscript revision.

S.F. contributed to data analysis and interpretation, and manuscript revision.

B.L. contributed to study conception and design, the acquisition and interpretation of data, as well as writing of this manuscript.

