## Supplemental Materials for "Sleep selectively and durably enhances real-world sequence memory"

### Supplementary Information for
Sleep selectively and durably enhances real-world sequence memory

Diamond, N.B*, Simpson, S.*, Baena Pérez, D., Murray, B., Fogel, S., & Levine, B.

* These authors contributed equally to the manuscript

Correspondence concerning this article should be addressed to Dr. Brian Levine


**This PDF file includes:**

Supporting Methods and Materials
Figures S1 to S6
Tables S1 to S5
SI References

### Supplementary Methods

#### Participants

##### *Study 1*

Fifty-seven healthy adults (14 males, 43 females; 18-35 years old, $M_{Age}$ = 22.74 years ± $SD_{Age}$ = 3.59 years) were recruited from the Baycrest participant registry and the community. All participants were naïve to the second floor of Baycrest, where the tour took place. Four participants were excluded: one for failing to complete one of the remote memory tests within the required time interval, one for technical issues in the headphones used for the tour audio guide, and two for test performance below 2.5 standard deviations from the sample mean on any of the four memory tests (averaged across featural and sequence items), which was suggestive of inattentive responding. After exclusions, data from 53 younger adults were included in analyses ($M_{Age}$ = 22.60 years ± $SD_{Age}$ = 3.23; $M_{Education}$ = 15.40 ± $SD_{Education}$ = 1.67).

##### *Study 2*

Ninety healthy adults (34 males, 56 females; 18 – 45 years old) were randomly assigned to either a Sleep group (*n* = 44, $M_{\text{Age}} \pm SD_{\text{Age}}$ = 24.09 ± 5.21 years; $M_{\text{Education}} \pm SD_{\text{Education}}$ = 15.66 ± 2.82 years) or a Wake group (*n* = 46, $M_{\text{Age}} \pm SD_{\text{Age}}$ = 24.57 ± 6.20 years; $M_{\text{Education}} \pm SD_{\text{Education}}$ = 15.43 ± 2.52 years). Participants underwent the same audio-guided walking tour as in Study 1 (a complete list of all audio tracks can be found on our OSF page: https://osf.io/bxm5w/). Five participants in the Wake group were excluded: three for failing to complete the second follow-up memory test before they went to sleep, one for later disclosing that they had been exposed to the second floor of Baycrest prior to the experiment, and one for taking a nap during the retention interval. One participant in the Sleep group who slept for only two hours was excluded. One participant in the Sleep group withdrew from the study. Four participants (three from the T1-T2 analysis, one from the T1-T4 analysis) were excluded for test performance below 2.5 standard deviations from the sample mean on any of the four memory tests (averaged across featural and sequence items). Finally, two participants failed to complete testing at T4 (these participants were retained for analyses involving T1-T2). As a result, 80 participants (*n* = 39 Wake, *n* = 41 Sleep) were included in the total sample, but only 77 (*n* = 38 Wake, *n* = 39 Sleep) were included in the analysis exploring memory change across T1-T4. The 56 (*n* = 33 Sleep, *n* = 23 Wake) participants who returned for T5 did not statistically differ in age ($t(54) = 0.24$, $p = .813$) or years of education ($t(52) = 0.62$, $p = .539$) from the 25 Study 2 participants who declined to participate in our long-term follow-up.

Participants in the Sleep and Wake groups were matched in terms of chronotype and self-reported sleep habits. Chronotype was assessed with the *Morningness-Eveningness Questionnaire Self-Assessment*(1), a 19-item survey that measures individual differences in human circadian rhythms. Participants' ratings on this instrument did not differ across the Wake and Sleep groups (*M*'s = 45.66 and 44.97, *SD'*s = 6.89 and 8.17 for the Wake and Sleep groups, respectively, $t(75) = 0.40$, $p = .693$). There were no significant group differences in self-reported sleep habits such as sleep quality, sleep disturbances, and use of sleep medication as rated on the *Pittsburgh Sleep Quality Index (*PSQI)(2) a nine-item instrument that assesses average sleep quality, patterns, and efficiency over the past month (M's = 9.95 and 9.28, SDs = 6.89 and 4.91 for the Wake and Sleep groups, respectively, $t(75) = 0.49$, $p = 0.627$). A global score of five or more indicates poor sleep quality. The higher the score, the worse the self-reported sleep quality (3).

***Study 3***

Eighteen additional participants were recruited to increase the sample size of the sleep group for testing individual differences, prior to halting of the study due to the onset of the COVID-19 pandemic. Of the 62 participants who completed PSG, 13 were excluded due to study withdrawal (1), technical issues of sweat sway affecting PSG recording (4), sleep efficiency scores below 60% (6), spindle counts more than 2.5 standard deviations below the group average the C4 channel (1), and overall memory performance more than 2.5 standard deviations below the group average at baseline (1). The final sample of 49 participants included 29 females and 20 males ($M_{Age}$ + $SD_{Age}$ = 24.29 ± 5.03 years); $M_{Education}$ + $SD_{Education}$ = 15.47 ± 2.94 years).

#### Polysomnography (PSG) Methods

The PSG montage included electrooculography (EOG), EEG, electromyography (EMG), and electrocardiographic (ECG) activity. Electrode placements followed the international 10-20 system and were referenced to the contralateral mastoid (A1 and A2). Vertical and horizontal eye movements were recorded via EOG electrodes that were placed on the left outer canthus (LOC) and right outer canthus (ROC). EEG scalp electrodes were placed at six different locations: frontal (F3, F4), central (C3, C4), and occipital (O1, O2) since these areas are ideal for measuring cortical neural activity such as spindles and slow waves (4). Two EMG electrodes were placed on the chin and the anterior tibial muscles to assist with AASM scoring. This allowed us to extract macrostructure measures of interest including sleep duration, efficiency, latency, and wake time after sleep onset. Data from each EEG channel was collected with a 256-Hz sampling rate, high- and low-pass filtered at 0.3 and 35 Hz, respectively, with a 60-Hz notch filter before being exported from the Compumedics system into European Data Format (EDF) files.

##### *Spindle Detection*

Spindle detection was carried out using a previously published method that was validated against both multiple expert scorers and crowd-sourced from a large sample of non-experts (5). Prior to automatic detection, movement and arousal artifact appearing in N2 and N3 periods were manually rejected by three expert raters from all participants. The spindle data were then extracted from movement artifact-free, N2 and N3 sleep epochs using the EEGlab plugin (6) “*detect_spindles2.2*” written for MATLAB R2019a (The MathWorks Inc., Natick, MA, United States). This detection method (5) uses a complex demodulation transformation of the EEG signal with a bandwidth of 5 Hz centred about a carrier frequency of 13.5 Hz (i.e., 11–16 Hz). This allowed us to extract the number, duration (s), amplitude (μV), and frequency (Hz) of spindles at each central derivation (C3 and C4). This spindle detection method was performed at C3 and C4 channels for slow (11–13.5 Hz), fast (13.5–16 Hz), and total bandwidth spindles (11–16 Hz). We analyzed total spindle counts averaged across both central channels; consideration of fast and slow spindles separately did not add new information to hypothesis-driven analyses. Due to an equipment issue, data for two participants was referenced to the same electrode (e.g., C3-A2 and C4-A2). In this event, we only selected data from the single contralaterally referenced electrode (e.g., C3-A2).

##### *Half-wave Detection*

The period amplitude analysis (PAA) method is adapted from well-established published methods  (7–9). The detection method requires narrow band-pass filtering to detect waves between 0.5 - 4 Hz, thus, derivations from F3 and F4 were band-pass filtered 0.46 Hz (64th-order Chebyshev type II high-pass filter, -80 dB stopband attenuation) and 4.30 Hz (32nd-order Chebyshev type II low-pass filter, -80 dB stopband attenuation) to achieve minimal attenuation in our 0.5 - 4 Hz band of interest and good attenuation at neighbouring frequencies. The filters were applied in the forward and reverse directions to achieve zero-phase distortion resulting in a doubling of the filter order. Half-waves were determined as negative or positive deflections between two consecutive zero crossings in the band-pass filtered signal. In line with scoring rules for slow waves, the peak-to-peak amplitude threshold of 75 μV (10) was applied to each half-wave and its neighbouring half-wave (i.e., half-wave pairs with < 75 μV peak-to-peak amplitude difference were excluded from further analysis). The duration of each half-wave that met the amplitude threshold was determined by the length of time (in seconds) between zero crossings. Here, we report the number of half-waves averaged over both channels as our proxy measure of SWA. Data from the two participants with referencing issues was treated in the same manner as detailed above.

##### *Supplemental Statistical Analyses*

Using the *performance* package in R, we confirmed that all of the effects in the linear mixed effect models were below a variance inflation factor of 5, indicating no issues with multicollinearity (11) (see tables below). For *a priori* *t*-tests, we used Welch's *t*-tests and reported Cohen's *d* effect sizes. Steiger's Z tests were also conducted to test statistical differences between correlations. Note that although we conducted our statistical tests on model fits, we visualized descriptive statistics (i.e., “raw” means from individual participant data) for ease of interpretability.

#### Supplementary Results

##### Time elapsed between encoding and follow-up tests

The follow-up tests (T2-T5) conducted outside of the laboratory were completed online, unsupervised. We confirmed that participants were compliant with the instructions for completing the follow-up tests within the requested time interval. In Study 2, there were no group (Wake vs. Sleep) differences in the time elapsed between tour encoding and the 12-hour (T1-T2: $t(75) = 1.39$, $p = .170$, $d = 0.316$), 1-week (T1- T3: $t(75) = 1.03$, $p = .307$, $d = 0.234$), 1-month ($t(75) = -0.28$, $p = .782$, $d = -0.063$), or 1-year ($t(49) = 1.04$, $p = .302$, $d = 0.294$) delays.

**Table S1.** Time elapsed between encoding and follow-up tests for participants in Studies 1 and 2 (standard deviation reported in parentheses).

| Study | Delay between encoding and T2 (hours)* | Delay between encoding and T3 (days) | Delay between encoding and T4 (days) | Delay between encoding and T5 (days)* |
|---|---|---|---|---|
| Study 1 | 24.52 (3.64) | 6.98 (.24) | 28.31 (1.72) | –- |
| Study 2 (Sleep group; n = 39) | 10.97 (0.26) | 7.05 (0.17) | 28.02 (0.10) | 366.64 (7.95) |
| Study 2 (Wake group; n = 38) | 11.24 (1.20) | 7.61 (3.40) | 28.01 (0.18) | 368.89 (7.32) |

* Subset of Study 2 participants who performed T5 (*n* = 23 Wake, *n* = 26 Sleep).

**Differential effects of sleep on sequence and featural memory not accounted for by psychometric characteristics**

Our four *a priori* test forms were constructed to be equivalent in terms of item characteristics and difficulty across forms and administration sessions. Moreover, there was no differential effect of item difficulty, validity (true/false status) or content repetition on sequence versus featural memory.

A 2 (Retrieval Type: detail vs. sequence) × 2 (item validity: true vs. false) ANOVA revealed neither effects of Retrieval Type ($F$(1, 269) = 2.67, $p$ = .103) nor item validity ($F$(1, 269) = .38, $p$ = .540) on overall item accuracy across participants (see Figure S1).

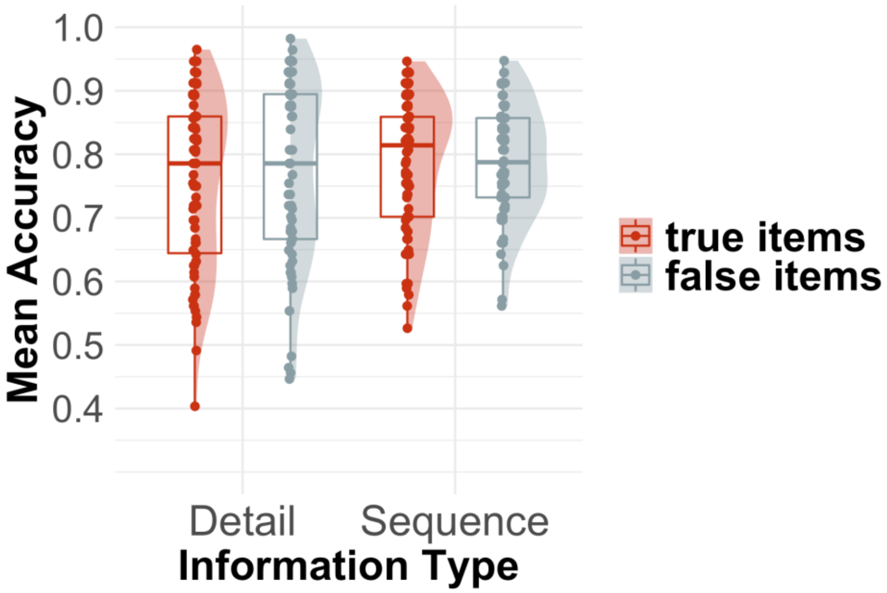


**Fig. S1***.* Average accuracy for items across participants in Study 1. Dots depict mean accuracy for individual test items, averaged across participants and timepoints. Shaded "half violin plots" and boxplots show the distribution of item-level accuracy across Retrieval Type and item validity.

We included item validity in our main model of sequence vs. featural memory change. Consistent with the above item-level analysis, at the participant level, item validity did not predict response accuracy ($\chi^2$(1) = .289, $p$ = .591) and did not interact with Time ($\chi^2$(1) = .463, $p$ = .496) nor Retrieval Type ($\chi^2$(1) = .017, $p$ = .896), nor was there a three-way interaction ($\chi^2(1) = .739$, $p = .390$). This means that the decline in detail memory over time, and the difference between featural and sequence memory change, were not driven differently by misses (responding ‘false’ to a true item) and false alarms (responding ‘true’ to a false item).

We considered that the contrasting forgetting curves for featural and sequence memory could be partly due to differential effects of repeated testing, rather than by time per se. Sequence statements, though trial-unique, comprise pairwise recombinations of a common set of tour events (e.g. “You encountered X before Y” and “You encountered Z before X”). Conversely, feature statements each referred to a specific item or tour feature, though for each target item in the tour we probed multiple features (e.g., its colour, size, etc.). If sequence memory benefited uniquely across the testing procedure via latent reactivation of intervening items (12), then sequence memory performance would increase across trials *within* a test, either in absolute terms or relative to featural memory performance. We did not find evidence for differential featural and sequence memory performance across trials within T1: there was neither a main effect of trial number ($\beta = 0.002$, $z = -.84$, $p = .40$) on memory accuracy across successive T1 trials, nor an interaction between trial number and retrieval type ($\beta = 0.01$, $z = 1.12$, $p = .234$), suggesting that sleep and/or time transform sequence versus featural memory change over and above repeated testing.

***Sequence Lag Effects***

As expected, sequence memory accuracy increased as the inter-item lag between items in question increased (main effects of lag in Study 1 and Study 2: $\chi^2(2) = 8.802$, $p = .012$, and $\chi^2(2) = 8.59$, $p = .014$) (see Figure S2). In Study 1, there was no significant effect of Time ($\chi^2(1) = .466$, $p = .495$), but in Study 2 there was ($\chi^2(1) = 12.59$, $p = < .001$); compared to T1, sequence memory across all lag levels was preserved after 12 hours ($z = 1.56$, $p = 0.238$) and 1 week ($z = 0.71$, $p = 0.476$), and only showed signs of decline after one month ($z = 4.41$, $p < 0.001$).

We considered the possibility that shorter lag pairs – requiring finer-grained spatiotemporal context retrieval – would decline more over time than coarser-grained far pairs, which could indicate a transformation from more specific to more gist-like memory for sequence information. We did not observe evidence for this effect: the Lag × Time interaction effect was neither significant in Study 1 ($\chi^2(2) = 1.74$, $p = 0.418$), nor in Study 2 ($\chi^2(2) = 1.79$, $p = 0.408$), nor did lag interact with Group ($\chi^2(2) = 2.55$, $p = 0.279$) or Group × Time in Study 2 ($\chi^2(2) = 2.08$, $p = 0.353$). These findings suggest that although finer-grained spatiotemporal context representations are more difficult to retrieve overall, they are not forgotten significantly faster than coarser-grained representations, nor are they differentially affected by sleep per se.

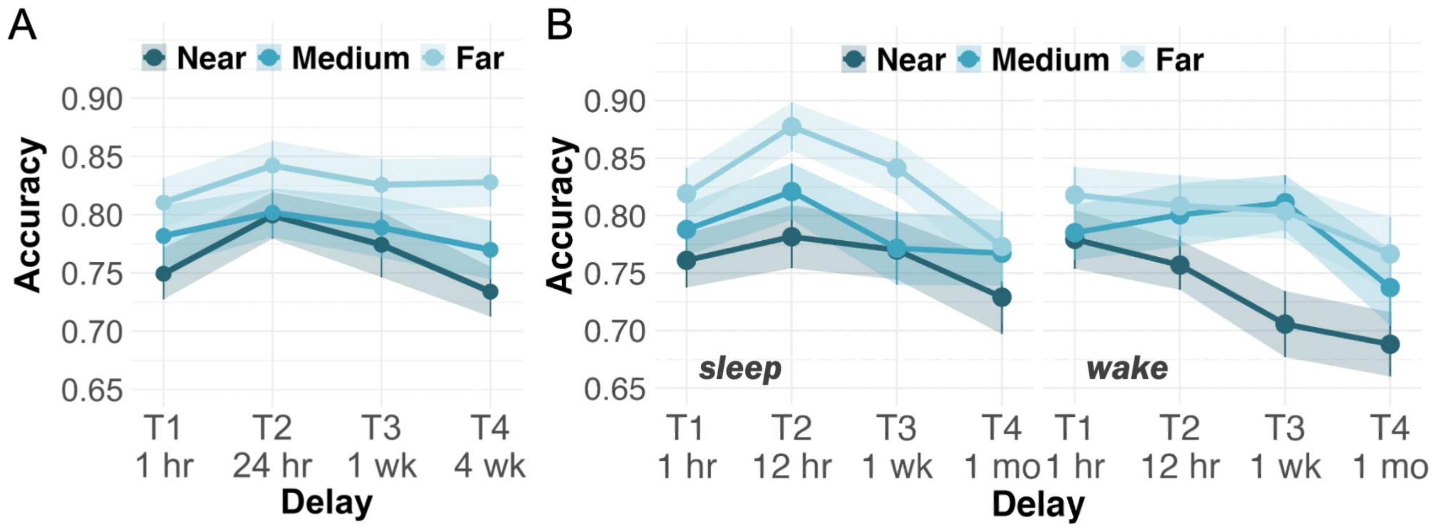


**Fig. S2.** Effects of inter-item lag on sequence memory accuracy in Study 1 (panel A), and Study 2 (panel B).

**Study 2: Individual Differences in Overnight Memory Change**

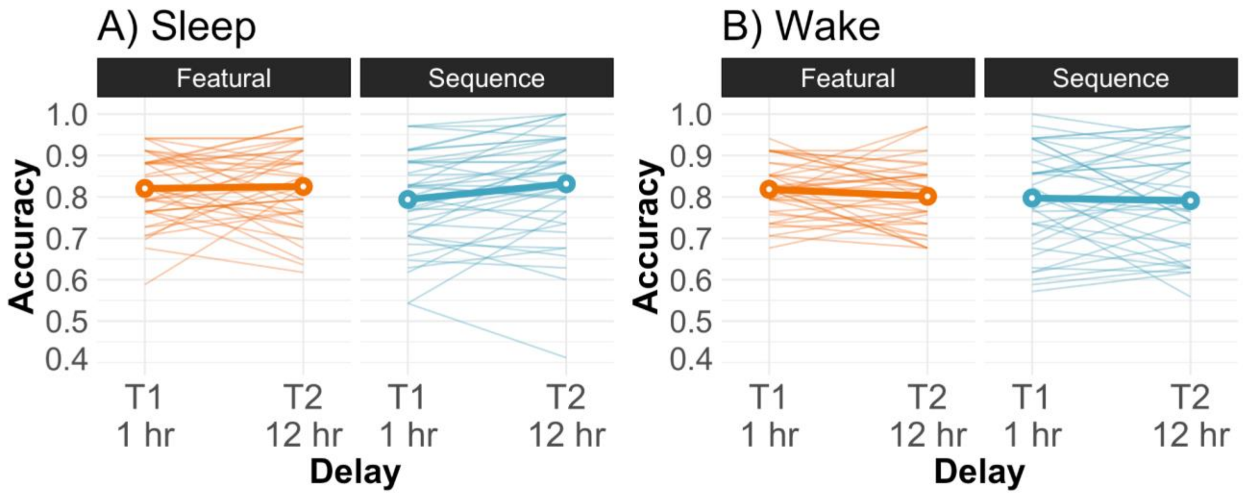


**Fig. S3.** Individual differences in overnight featural (orange) and sequence (blue) memory (at 1 hr and 12 hr delays) either after a night of sleep (panel A) or an equivalent period of wakefulness (panel B) for participants in Study 2. Thick circles with white fill and thick solid lines depict group averages; each thin line depicts an individual participant.

**Study 3 Results**

**Table S2.** Descriptive statistics of polysomnography and sleep survey data from the larger sleep sample (*N* = 49). Standard deviations are included in parentheses.

| | Mean (SD) | Minimum | Maximum |
|---|---|---|---|
| Total sleep time (min) | 420.14 (35.08) | 312.5 | 469 |
| Sleep efficiency (%) | 87.64 (7.09) | 68.9 | 96.8 |
| N1 duration (min) | 41.14 (19.57) | 15 | 103 |
| N2 duration (min) | 224.76 (38.10) | 147.5 | 302 |
| SWS duration (min) | 87.40 (29.96) | 24 | 166.5 |
| REM duration (min) | 65.22 (20.18) | 30 | 118.5 |

| | | | |
|---|---|---|---|
| Non-REM duration (min) | 353.30 (35.31) | 259.5 | 396.5 |
| Average spindle count (raw value) | 1505.02 (258.65) | 1024 | 1984.5 |
| Average half-wave count (raw value) | 10634.79 (3627.85) | 2322 | 18579 |

**Table S3.** Effects from linear mixed effects model assessing relationship of sleep macrostructure variables to memory performance on T1 and T2. The dependent variable was each participant's average memory score at each timepoint split by featural vs. sequence (where memory change is assessed via interactions with Time); therefore, only effects involving the Time factor (i.e., T1, T2; immediate vs 12-hr testing) are listed. The interaction between Time and Total Sleep Time (TST) was included as a covariate of non-interest. Asterisks indicate statistically significant effects. VIF = Variance Inflation Factor. Df = degrees of freedom; Df res = residual degrees of freedom

| Effect | *F* value | Df | Df res | *p* value | VIF |
|---|---|---|---|---|---|
| Time × Retrieval type | 1.85 | 1 | 131 | .176 | 2.00 |
| Time × N1 duration | 1.09 | 1 | 131 | .298 | 3.77 |
| Time × N2 duration | 1.94 | 1 | 131 | .167 | 8.67 |
| Time × SWS duration | 4.51 | 1 | 131 | .036 * | 7.25 |
| Time x REM duration | 3.84 | 1 | 131 | .052 | 2.95 |
| Time x TST | 0.82 | 1 | 131 | .368 | 7.69 |
| Time × Retrieval type × N1 duration | 0.09 | 1 | 131 | .766 | 3.14 |
| Time × Retrieval type × N2 duration | 0.21 | 1 | 131 | .643 | 2.66 |
| Time × Retrieval type x SWS duration | 0.01 | 1 | 131 | .934 | 3.34 |

| | | | | | |
|---|---|---|---|---|---|
| Time × Retrieval type × REM duration | 0.07 | 1 | 131 | .787 | 2.17 |

**Table S4.** Effects of linear mixed effects model assessing the relationships of sleep spindles and half-waves to memory performance (in separate models). Only effects involving the Time factor (i.e., T1, T2; immediate vs 12-hr testing) are listed. Asterisks indicate statistically significant effects. VIF = Variance Inflation Factor. Df = degrees of freedom; Df res = residual degrees of freedom.

| Effect | *F* value | Df | Df res | *p* value | VIF |
|---|---|---|---|---|---|
| **Spindles** | | | | | |
| Time × Retrieval Type | 1.92 | 1 | 141 | .168 | 2.00 |
| Time × Average Spindle Count | 8.66 | 1 | 141 | .004 * | 1.18 |
| Retrieval Type × Average Spindle Count | 0.01 | 1 | 141 | .959 | 2.00 |
| Time x Retrieval Type × Average Spindle Count | 0.06 | 1 | 141 | .812 | 2.00 |
| **Half-waves** | | | | | |
| Time × Retrieval Type | 1.91 | 1 | 141 | .169 | 2.00 |
| Time × Average Half-wave Count | 7.43 | 1 | 141 | .007 * | 1.18 |
| Time x Retrieval Type × Average Half-wave Count | 0.06 | 1 | 141 | .806 | 2.00 |

**Table S5.** Effects of linear mixed effects model assessing the relationships of coupled vs. uncoupled spindles to memory performance. Only effects involving the Time factor (i.e., T1, T2; immediate vs 12-hr testing) are listed. Asterisks indicate statistically significant effects. VIF = Variance Inflation Factor. Df = degrees of freedom; Df res = residual degrees of freedom.

| Effect | *F* value | Df | Df res | *p* value | VIF |
|---|---|---|---|---|---|

| | | | | | |
|---|---|---|---|---|---|
| Time × Retrieval Type | 1.90 | 1 | 138 | 0.170 | 2.00 |
| Time × Average Coupled Spindles | 9.32 | 1 | 138 | 0.003 * | 1.21 |
| Time × Average Uncoupled Spindles | 0.07 | 1 | 138 | 0.792 | 1.21 |
| Time × Retrieval Type × Average Coupled Spindles | 0.01 | 1 | 138 | 0.909 | 2.02 |
| Time × Retrieval Type × Average Uncoupled Spindles | 0.22 | 1 | 138 | 0.640 | 2.02 |

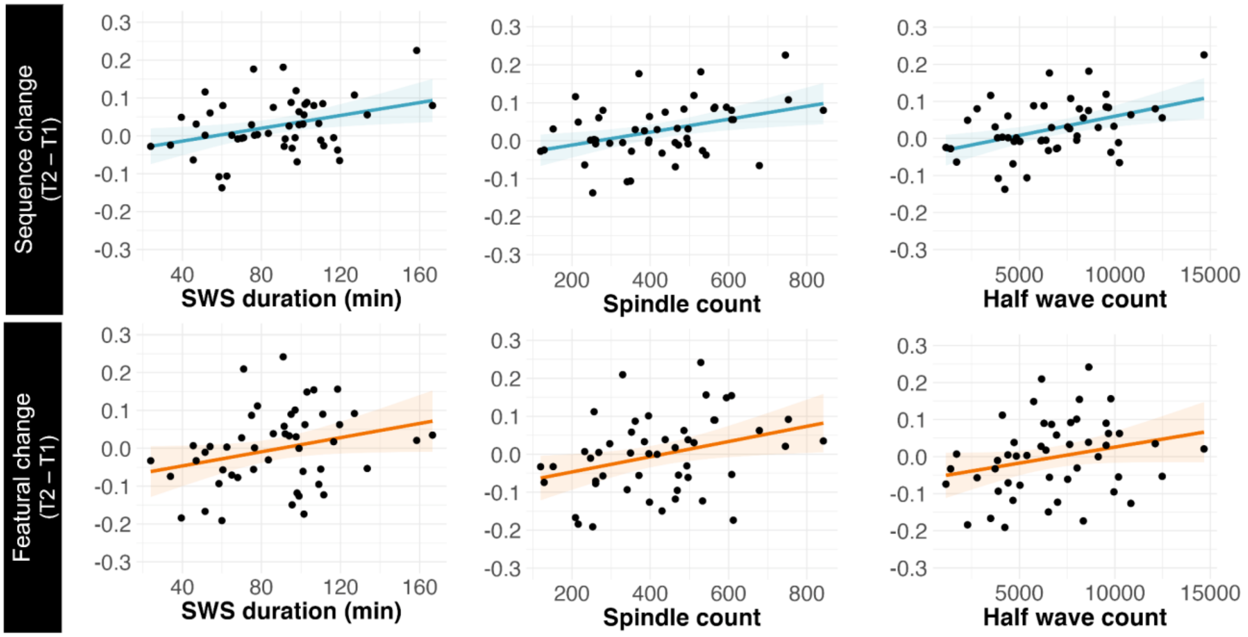


**Fig. S4.** Correlation between SWS duration, spindles, and half-waves and overnight (T1 to T2) change in memory for sequence (top panel) and features (bottom panel). Sequence memory is positively correlated with SWS ($r(47) = 0.35$, $p = .015$), spindle count ($r(47) = 0.38$, $p = .006$), and half-wave ($r(47) = 0.42$, $p = .002$) count. Featural memory is positively related to SWS ($r(47) = 0.28$, $p = .055$), spindle count ($r(47) = 0.33$, $p = .022$) and half-wave count ($r(47) = 0.25$, $p = .077$). Overall memory is positively correlated with SWS duration ($r(47) = 0.38$, $p = .007$), spindle count ($r(47) = 0.44$, $p = .002$), and half-wave count ($r(47) = 0.41$, $p = .004$) (not pictured here).

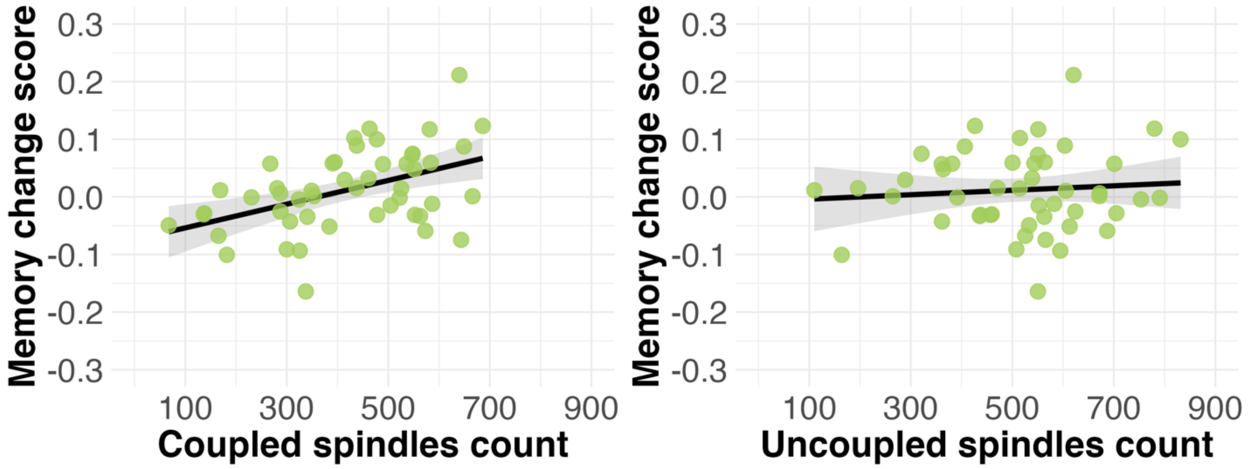


**Fig. S5.** Correlation between coupled vs. uncoupled spindles in SWS and overnight (T1 to T2) overall memory change. The correlations for coupled spindles ($r(47) = 0.46$, $p < .001$) and uncoupled spindles ($r(47) = 0.09$, $p = .545$) are statistically different from each other, Steiger's $z = 2.11$, $p = .038$.

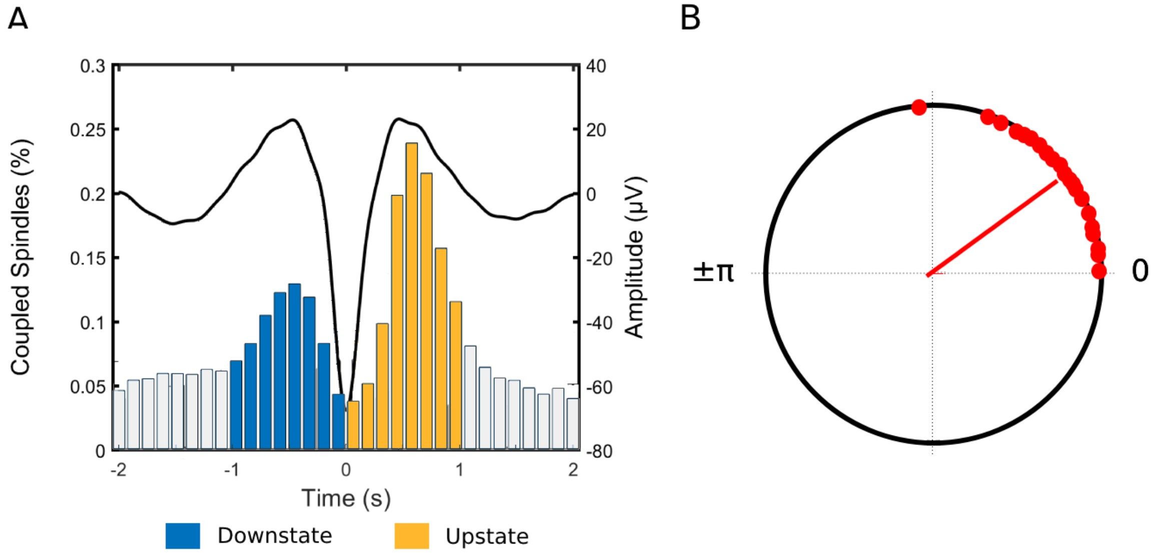


**Fig. S6. A)** Coupled slow wave-spindle histogram for all participants. Each bar represents the number of coupled spindles detected in an interval of 125 milliseconds divided by the total number of spindles. Average slow wave oscillation for all participants is superimposed in black. **B)** Circular plot of preferred phase for each individual (slow wave phase at spindle amplitude maximum). Red dots denote an individual preferred phase ($0^{o}$ slow wave down-to-up state, ± $180^{o}$ slow wave up-to-down state). The direction of the line indicates the preferred direction of the grand average. Most individuals exhibit spindles adjacent to, or immediately preceding, the positive slow wave peak at $0^{o}$.
